# Frequency Shift Imaging Using an Adiabatic SLR Self-Refocused Pulse for Positive-Contrast SPION Imaging at 7T

**DOI:** 10.64898/2026.09.27.754833

**Authors:** Akbar Alipour, Madison Chiu, Judy Alper, Hadrien Dyvorne, Gaura Verma, Shams Rashid, Abraham J. P. Teunissen, Zahi Fayad, Priti Balchandani

**Author notes:** Corresponding Author: Akbar Alipour, MSc, PhD, JFISMRM Assistant Professor, BioMedical Engineering and Imaging Institute (BMEII) Department of Diagnostic, Molecular and Interventional Radiology Icahn School of Medicine at Mount Sinai 3 East 101st Street, 7th Floor, Room 708, New York, 10029.

## Abstract

**Purpose:** Superparamagnetic iron oxide nanoparticles (SPION) are widely used as MRI contrast agents for cell tracking and molecular imaging. However, their detection using conventional T_2_/T_2_* weighted and susceptibility-weighted MRI relies on negative contrast (signal voids), which lacks specificity because similar hypointense signals can arise from other sources of magnetic susceptibility. This study introduces Frequency Shift Imaging (FSI), a positive-contrast technique based on an adiabatic Shinnar-Le Roux (SLR) self-refocused pulse that addresses the limited specificity of conventional SPION imaging.

**Methods:** An adiabatic self-refocused RF pulse was designed using the SLR algorithm for frequency-selective positive-contrast imaging. The optimized pulse had a spectral bandwidth of 370 Hz, a peak RF amplitude of 14 μT, and a duration of 15 ms. FSI data were acquired on a 7T MRI scanner. Phantom experiments used agarose-embedded 30 nm cell-labeled SPION at 0%, 25%, 75%, and 100% concentrations (pure). In vivo validation was performed following intramuscular SPION injection into the hind limb of a mouse. FSI images were compared with conventional acquisitions.

**Results:** Quantitative phantom analysis showed normalized FSI signal intensities of 25.02% and 75.16% for samples containing 25% and 75% labeled cells, respectively, demonstrating an approximately proportional relationship between FSI signal and labeled-cell concentration. In vivo, FSI generated localized hyperintense positive contrast at the SPIO injection site, providing improved conspicuity and localization compared with the corresponding negative-contrast signal void.

**Discussion and Conclusion:** FSI enables robust, B_1_-insensitive, positive-contrast SPION imaging at 7T with reduced echo time and SAR compared to paired adiabatic approaches, supporting its potential for quantitative cell tracking and molecular imaging applications.

## Introduction

Magnetic resonance imaging (MRI) has undergone substantial evolution with the development of advanced contrast agents, expanding its role beyond anatomical imaging into molecular and cellular applications [1, 2]. These advances have enabled noninvasive monitoring of molecular pathways, in vivo cell tracking, and assessment of targeted therapeutic interventions [3, 4]. Among available contrast agents, superparamagnetic iron oxide nanoparticles (SPION) have attracted considerable interest due to their strong magnetic susceptibility, high biocompatibility, and ease of functionalization for targeted labeling [5, 6]. Consequently, SPION are widely utilized in both preclinical and clinical settings to investigate cellular migration, tumor progression, and inflammatory processes [7–9]. The versatility of SPION has further expanded with the development of ultrasmall formulations capable of T_1_-weighted contrast enhancement, offering a potentially safer alternative to gadolinium-based agents [10, 11].

SPION primarily act by inducing local magnetic field inhomogeneities, which lead to rapid dephasing of nearby water proton spins [12, 13]. This effect results in shortening of the transverse relaxation times (T_2_ and T_2*_), producing hypointense regions, or signal voids, on T_2_- and T_2*_- weighted images[14]. While this negative-contrast mechanism is highly sensitive, it has limited specificity, as signal voids can be confounded by other sources of susceptibility variation, including air-tissue interfaces, hemorrhage, and imaging artifacts. Negative contrast also complicates quantitative assessment because the degree of signal loss depends on an underlying baseline signal that cannot be directly measured within the signal void [15, 16]. Additionally, negative contrast is susceptible to partial-volume effects, requiring voxel sizes significantly smaller than the induced susceptibility region for accurate detection [17]. Paired pre- and post-contrast MRI scans are therefore recommended to improve the accuracy of image interpretation and artifact elimination [^18^]. Although positive contrast with SPIONs can be achieved using T_1_- weighted imaging, this approach is generally restricted to very small nanoparticles and may be limited by dominant (T_2_/T_2*_) effects that obscure T_1_ shortening. From a quantitative perspective, positive-contrast approaches may offer an advantage over negative contrast because the measured hyperintense signal can be directly related to the SPION-associated response, whereas quantification from a signal void is more challenging because the underlying baseline signal within the void cannot be directly measured [17, 19–21].

To address these limitations, several positive-contrast imaging techniques have been developed to convert SPION-induced signal voids into hyperintense signals. These methods include Gradient Echo Acquisition for Superparamagnetic Particles with Positive Contrast (GRASP), which suppresses background signal using tailored gradients; Inversion-Recovery with ON-resonant water suppression (IRON), which highlights off-resonant spins; Fast Low-Angle Positive Contrast Steady-State (FLAPS); and spectrally selective excitation approaches that directly target off-resonant water protons near SPION [22–25]. Among these, selective excitation techniques are particularly promising because they exploit the frequency shifts induced by SPION- generated field perturbations [26, 27]. Early implementations employed paired excitation and refocusing pulses tuned to specific frequency offsets, enabling positive contrast that scales linearly with SPION concentration [27]. However, these methods often lacked slice selectivity and required long echo times (TE), increasing signal loss due to diffusion [26]. To address these challenges, Balchandani et al. introduced the self-refocused spatial-spectral (SR-SPSP) pulse, designed using the Shinnar-Le Roux (SLR) algorithm, which combines excitation and refocusing into a single pulse to reduce TE and improve spatial localization [27].

Ultra-high-field MRI at 7T offers increased signal-to-noise ratio (SNR) and enhanced sensitivity to magnetic susceptibility effects, which can improve the detection of SPIO-induced field perturbations and make 7T particularly attractive for SPIO-based imaging [28, 29]. In addition, the high spatial resolution and anatomical detail achievable with structural T_1_- and T_2_- weighted imaging at 7T can facilitate accurate localization of SPIO-associated signals when SPIO-based images are co-registered with the underlying anatomical images. However, these advantages are accompanied by increased B_0_ and B_1_ inhomogeneities and greater radiofrequency (RF) power deposition, which can compromise image uniformity and limit pulse-sequence performance. These competing advantages and technical challenges motivate the development of robust, B_1_-insensitive positive-contrast approaches specifically suited for SPIO detection at 7T. At 7T, pronounced B_1_ inhomogeneity can lead to spatially varying flip angles and nonuniform spin-echo (SE) signals, motivating the use of adiabatic RF pulses for improved B_1_ robustness [^3^0-3^4^]. However, conventional adiabatic refocusing schemes often require additional RF pulses to compensate for nonlinear phase, increasing TE and specific absorption rate (SAR). A self-refocused adiabatic pulse previously developed by our group addresses these limitations at 7T MRI by integrating excitation and refocusing into a single RF pulse while maintaining adiabatic robustness, thereby reducing TE and RF power deposition [30].

In this work, we introduce a Frequency Shift Imaging (FSI) technique that leverages the SLR framework to generate a matched-phase SE adiabatic 90°-180° pulse pair. The narrow spectral bandwidth of the FSI pulse enables highly selective excitation of off-resonant spins in the vicinity of SPIOs, while the adiabatic design ensures robust performance in the presence of B_1_ inhomogeneity at 7T [35–37]. This approach enables efficient positive-contrast imaging of SPIOs at ultra-high field strengths, which we validate through both phantom and in vivo experiments.

## 2. Methods

### 2.1 Filter Design

The FSI pulse was designed using the adiabatic SLR framework. Figure 1 illustrates the pulse- design algorithm and the main steps involved in generating the RF pulse. In the SLR framework, RF pulse design is represented by two polynomials, A_n_(z) and B_n_(z), which are used as inputs to the inverse SLR transform to generate the corresponding RF pulse waveform[32, 33]. Because these polynomials can be represented as finite impulse response (FIR) filters, conventional digital filter design techniques can be used to specify their spectral characteristics. Typically, B_n_(z) is first designed to generate the desired frequency response, after which the corresponding minimum- phase A_n_(z) is derived [^32^, ^33^]. The resulting polynomial pair defines the spin rotation and provides a minimum-energy RF pulse with the prescribed spectral profile.

**Figure 1.**
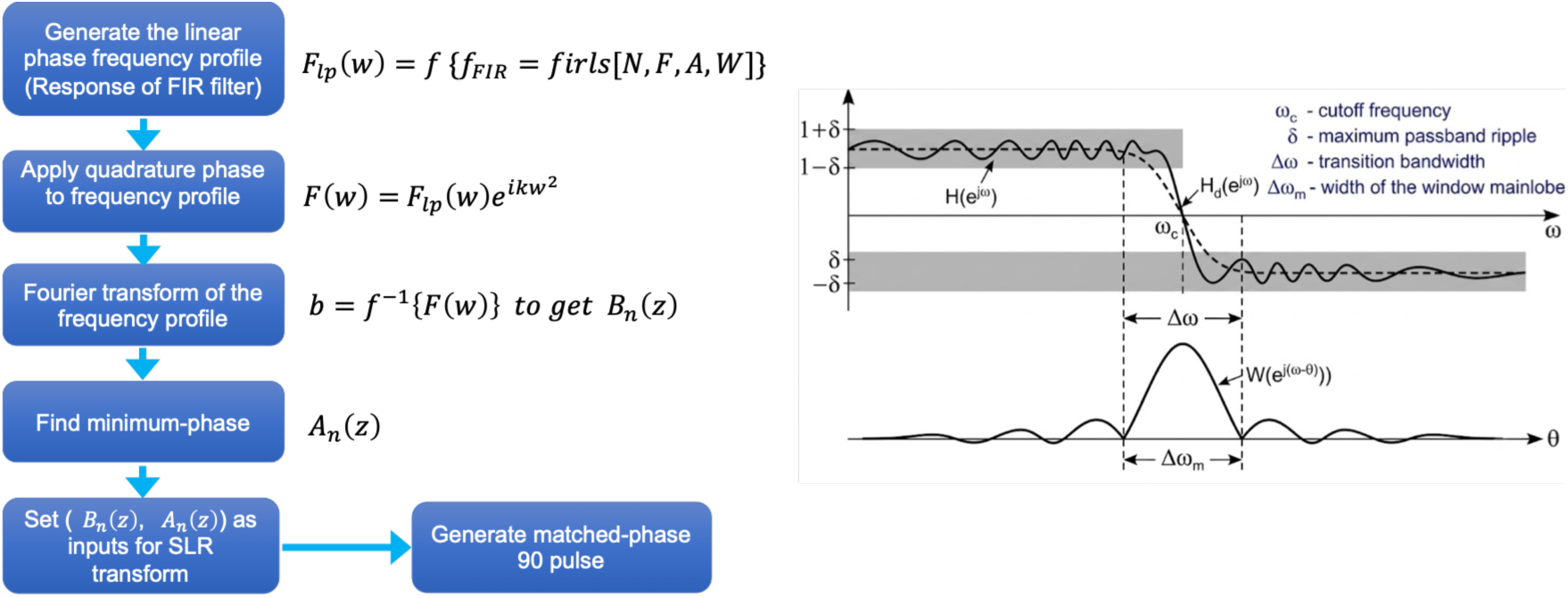
Design of the FSI RF pulse. The flowchart on the right illustrates the adiabatic SLR pulse-design process and generation of the self-refocused pulse. The left panel shows the target spectral filter response and key parameters used to define the frequency-selective RF pulse.

The first step in the FSI pulse design pipeline is the construction of a narrow-band FIR lowpass filter whose frequency-domain profile serves as the beta polynomial envelope for the SLR algorithm. The filter specification defines the allowable magnetization response in three distinct frequency regions: the passband, the transition band, and the stopband. These regions and their corresponding tolerance constraints are illustrated in Figure 1, which shows the frequency response *H*(*e*^j^*^ω^*) of the designed filter alongside the ideal desired response *H_d_*(*e*^j^*^ω^*). The response must remain entirely within the unshaded regions.

The cutoff frequency *ω_c_* defines the nominal boundary between the passband and the stopband, located at the midpoint of the transition band. The position of *ω_c_* is indicated by the vertical dashed line in Figure 1 and is marked on the frequency axis below the plot. The ripple parameter *δ* specifies the maximum allowable deviation of the filter magnitude response from its ideal value within both the passband and the stopband. In the passband, the response is required to remain within the tolerance band about the ideal unity gain, as shown by the upper and lower shaded forbidden zones on the left side of Figure 1. The transition bandwidth *Δω* is the frequency interval separating the end of the passband specification from the start of the stopband specification. Within this interval, no constraint is imposed on the filter magnitude; the response is free to transition from the passband gain of approximately 1 to the stopband gain of approximately 0. The transition bandwidth is indicated in Figure 1 by the pair of arrows straddling the dashed *ω_c_* line. The main-lobe width *Δωₘ* characterizes the frequency-domain spread of the applied window function *W*(*e*^j(*ω*–*θ*)^) shown in the lower left panel of Figure 1. In windowed FIR filter design, convolving the ideal brick-wall frequency response with the window spectrum introduces a transition region whose width is directly proportional to *Δωₘ*. For a rectangular window (no windowing), the main-lobe is maximally narrow, but the sidelobes are high, producing the Gibbs oscillations visible as the ripple of the *H*(*e*^j*ω*^) curve in Figure 1. The relationship Δω ≈ Δωₘ holds approximately for windowed least-squares designs, linking the achievable transition bandwidth directly to the main-lobe width and, therefore, to the filter order.

Following the approach previously described by our group and others, the target frequency response of the adiabatic 180° pulse was defined using a least-squares linear-phase finite impulse response (FIR) filter implemented with the *firls* function, *f_FIR_* = *firls* (*N*, *F*, *A*, *W*) in MATLAB (MathWorks, Natick, MA) [32–34]. Where *N* is the number of coefficients, F= (1⁄*π*)[0 *ω_p_ ω_p_ π*], is a vector of frequency band edges given in the range [0, *π*] but normalized to [0,1], *A* is a vector that specifies the desired amplitude of the frequency response of the filter, in this case, is set for a lowpass filter, and, *W* = 7*δ_p_ δ_p_*9, contains the relative ripple amplitudes in the passband and stopband given by *δ_p_* and *δ_p_*, respectively that controls the relative passband and stopband ripple amplitudes. In our design, *N* =255, F= (1⁄*π*)[0 0.007*π* 0.01 *π*], A= [1 1 0 0], and *W* = [0.1/8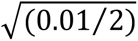0.01]. Following filter design, the impulse response was zero-padded to 4,000 samples in the frequency domain to obtain a high-resolution spectral profile for SLR processing. The resulting magnitude spectrum was then modulated by a quadratic phase function of the form *e*^j(*kω*2)^, with *k* = (3000⁄*π*^2^), to impose the frequency sweep necessary for adiabatic pulse behavior. This quadratic phase profile was applied via the *filter_pm* function, yielding the complex-valued frequency profile *F*(*ω*) = @*F_lp_*(*ω*)@*e*^j(^*^kω^* ^)^. The inverse Fourier transform of this profile produced the complex beta polynomial coefficients subsequently used in the SLR inversion.

The left panel of Figure 2 presents the magnitude responses on a logarithmic (dB) scale, zoomed to the transition region near the passband edge. The filter (orange) exhibited a stopband attenuation of only −26.7 dB, with sidelobe structure visible throughout the stopband. The optimized filter (blue) achieved −69.2 dB of stopband attenuation, a 42.5 dB improvement with a markedly cleaner spectral roll-off in the transition band. These improvements arose directly from the higher filter order, which provides finer frequency resolution, combined with the strongly asymmetric weight assignment that directed the optimization to prioritize passband flatness over stopband suppression. The right panel of Figure 2 displays the magnitude responses on a linear scale, restricted to the passband and lower transition region. The original design exhibited a passband ripple of ±0.088 (8.8%), reflecting the Gibbs phenomenon inherent to the near-rectangular effective window produced by the nearly equal weighting scheme. The optimized design reduced the passband ripple to ±0.00001, an 8,800-fold improvement yielding a visually flat passband response across the entire specified band (green shading, 0 to 0.004 × *π* rad/sample). This near-unity passband gain was essential for producing a flat-top FSI difference spectrum, as any passband ripple in the filter translates directly into amplitude modulation of the magnetization profile through the SLR transformation. The improvement in stopband attenuation (red shading) further reduced spectral leakage that would otherwise contaminate the FSI signal with off-resonance contributions.

**Figure 2.**
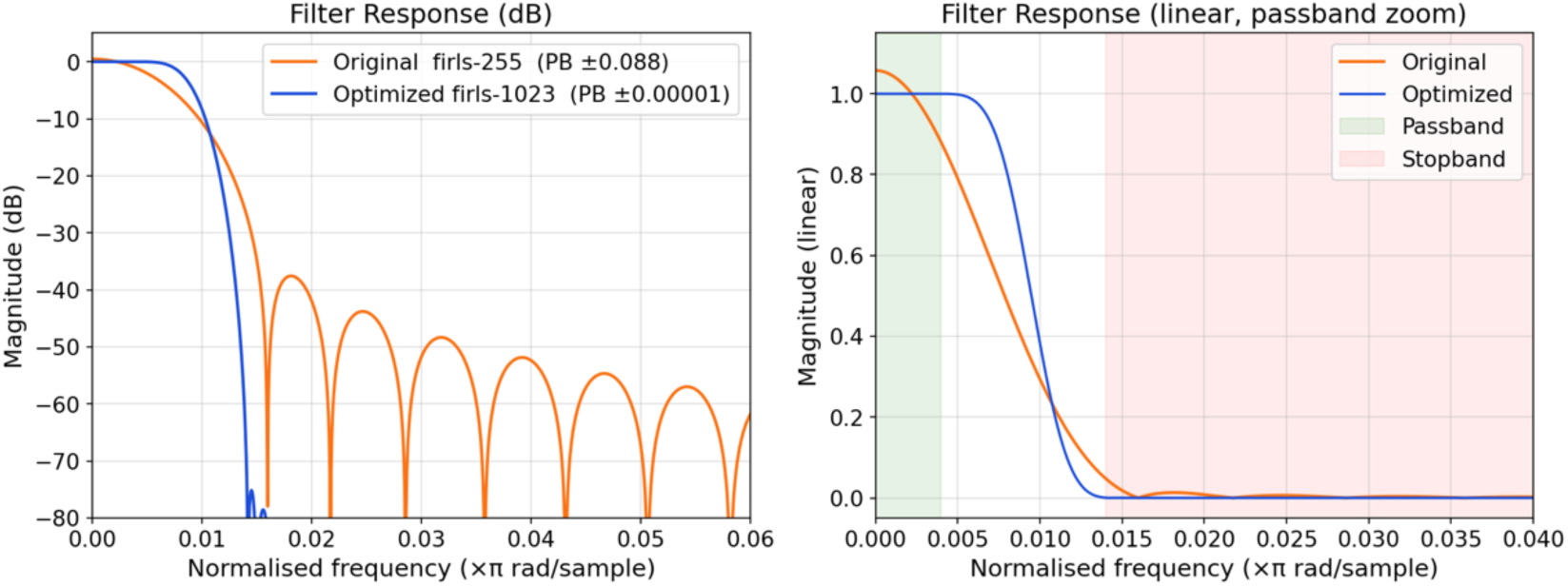
Comparison of the original and optimized FIR filter responses. Left: Log-scale magnitude response highlighting improved stopband attenuation and transition-band performance. Right: Linear-scale passband response demonstrating substantially reduced passband ripple in the optimized filter.

### 2.2. FSI RF Pulse Design

The resulting frequency profile of the filter was transformed into the B_180_ (z) polynomial using a Fourier transform, and the corresponding minimum-phase A_180_ (z) polynomial was subsequently derived. Both polynomials were then incorporated into the inverse SLR transform to generate the 180° RF pulse waveform. Figure 3a illustrates the intermediate steps used to generate the desired spectral response of the RF pulse. The magnitude and phase of the B_180_ (z) polynomial coefficients are shown after applying the quadratic phase to the FIR filter generated using the MATLAB firls function. The corresponding magnitude and phase of the Fourier-transformed B_180_ (z) polynomial demonstrate the resulting frequency-domain response, with the magnitude profile in Figure 3b defining the target spectral profile of the RF pulse. The initial pulse design had a spectral bandwidth of 370 Hz, a duration of 8 ms, and a peak RF amplitude of 14 μT. This narrow passband is essential for FSI, where only a chemically specific, frequency-shifted magnetization component must be selectively excited while all other spectral components remain unperturbed.

**Figure 3.**
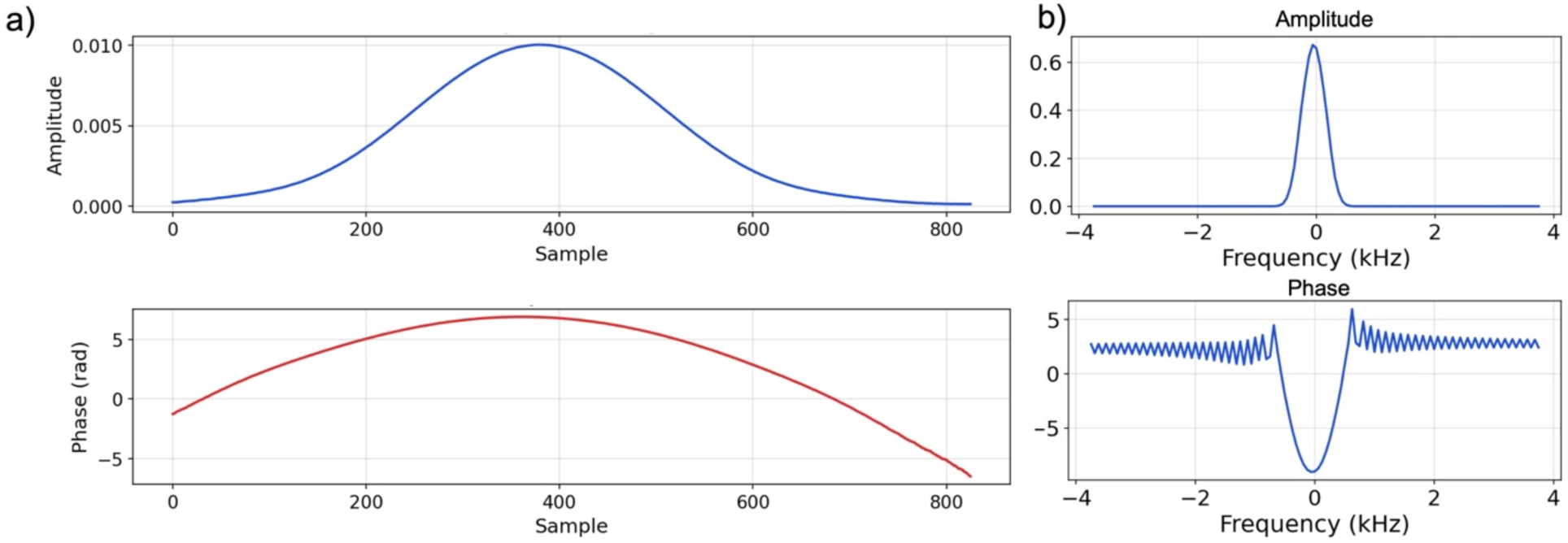
a) Magnitude and phase of the B polynomial coefficients after applying quadratic phase to the FIR filter. b) Magnitude and phase of the corresponding Fourier-transformed B polynomial, representing the desired spectral profile.

The designed adiabatic 180° pulse was subsequently used to construct a phase-matched 90°- 180° pulse pair and the corresponding self-refocused adiabatic FSI pulse. This implementation followed the framework described in our previous works, in which the spectral phase of the 90° excitation pulse is matched to that of the adiabatic 180° pulse to compensate for its nonlinear phase [30, 36]. The two-pulse configuration is then represented by a single set of SLR polynomials designed to produce transverse magnetization equivalent to that obtained with the separate 90° excitation and 180° refocusing pulses.

The B_180_ (z) polynomial of the adiabatic refocusing pulse was used to derive the B_90_ (z) polynomial for the phase-matched 90° excitation pulse according to Eq. 1.

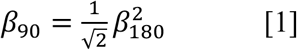

After obtaining B_90_ (z) the corresponding minimum-phase A_90_ (z) polynomial was calculated using the established SLR formulation [^16^]. Together with the previously determined A_180_ (z) and B_180_ (z) polynomials, these terms define the phase-matched adiabatic 90° excitation and 180° refocusing pulses. The RF amplitude and phase profiles of the 180° and 90° pulses are presented in Figure 4a and Figure 4b, respectively. The nonlinear spectral phases of the two pulses are matched such that their combined application produces a linear-phase SE response without requiring an additional 180° refocusing pulse.

**Figure 4.**
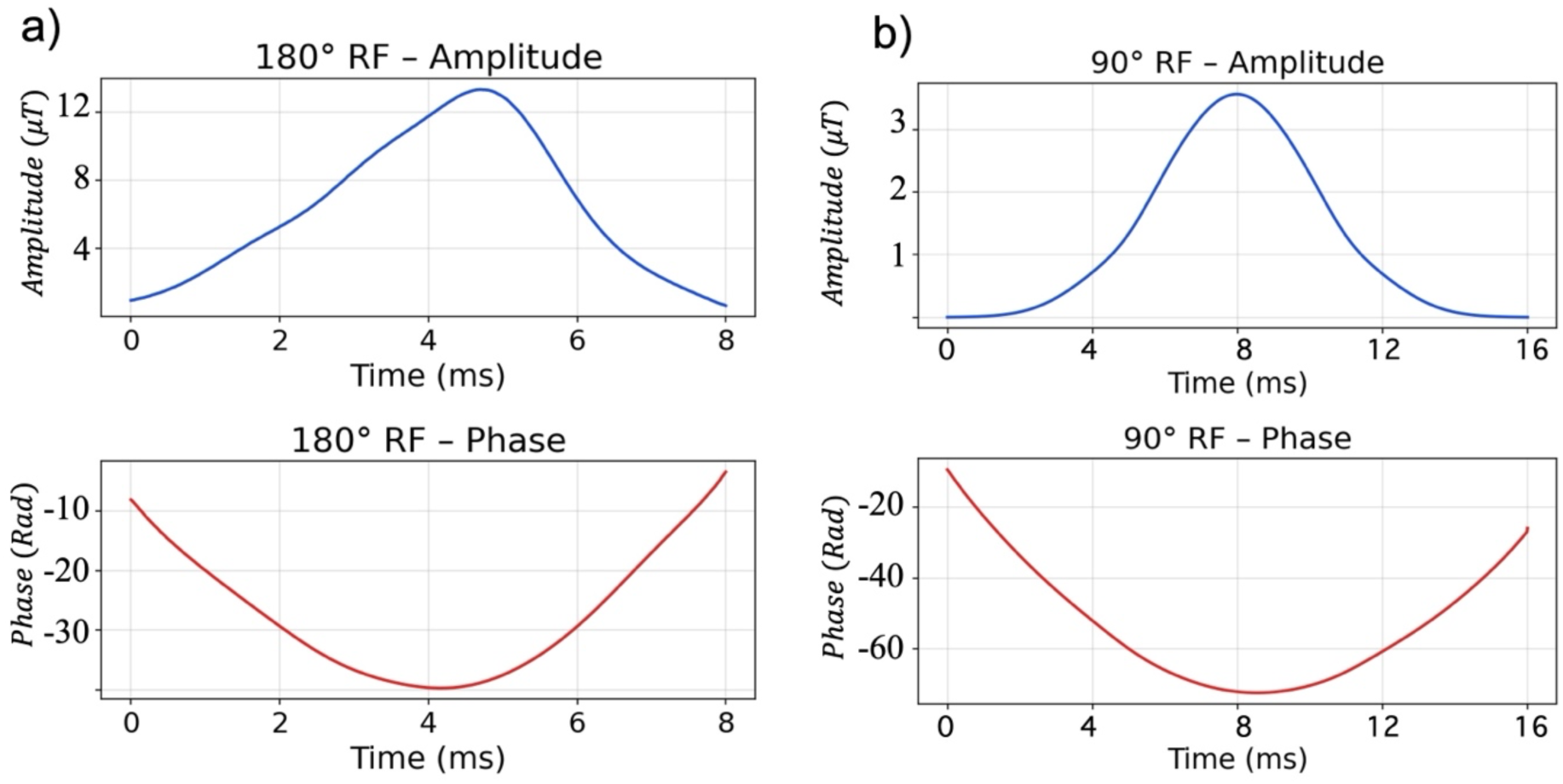
RF waveforms of the phase-matched pulse pair. a) Magnitude and phase of the adiabatic 180° pulse. b) Magnitude and phase of the corresponding phase-matched 90° pulse.

To further shorten the minimum TE, the phase-matched 90°-180° pair was reformulated as a single self-refocused adiabatic (SRA) pulse following the approach described in our previous work [^27^]. The B_90_ (z) and B_180_ (z) polynomials were used to derive the SRA polynomial pair, A_sr_ (z) and B_sr_ (z), such that the resulting pulse generates transverse magnetization equivalent to that produced by the separate excitation and refocusing pulses. If *α* and *β* denote the frequency-domain profiles associated with the SLR polynomials A_n_ (z) and B_n_ (z), respectively, the corresponding profiles *α_sr_* and *β_sr_* for the self-refocused pulse are defined according to Eq. 2.

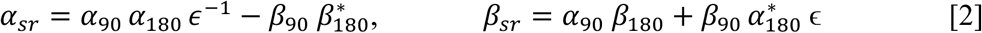

Here, *ε* = *e^iω^*^(Δ*T*)^, *ω* denotes the spectral frequency and Δ*T* represents the interval from the end of the self-refocused pulse to the formation of the spin echo. Incorporating the phase-matched 90° excitation and adiabatic 180° refocusing components within a single self-refocused pulse reduces the temporal separation between excitation and refocusing, thereby allowing a shorter minimum TE than can be achieved when the two RF pulses are applied separately in SE sequences. Figure 5 shows the amplitude and phase of the self-refocused pulse, which combines the 90° excitation and 180° refocusing pulses into a single pulse.

**Figure 5.**
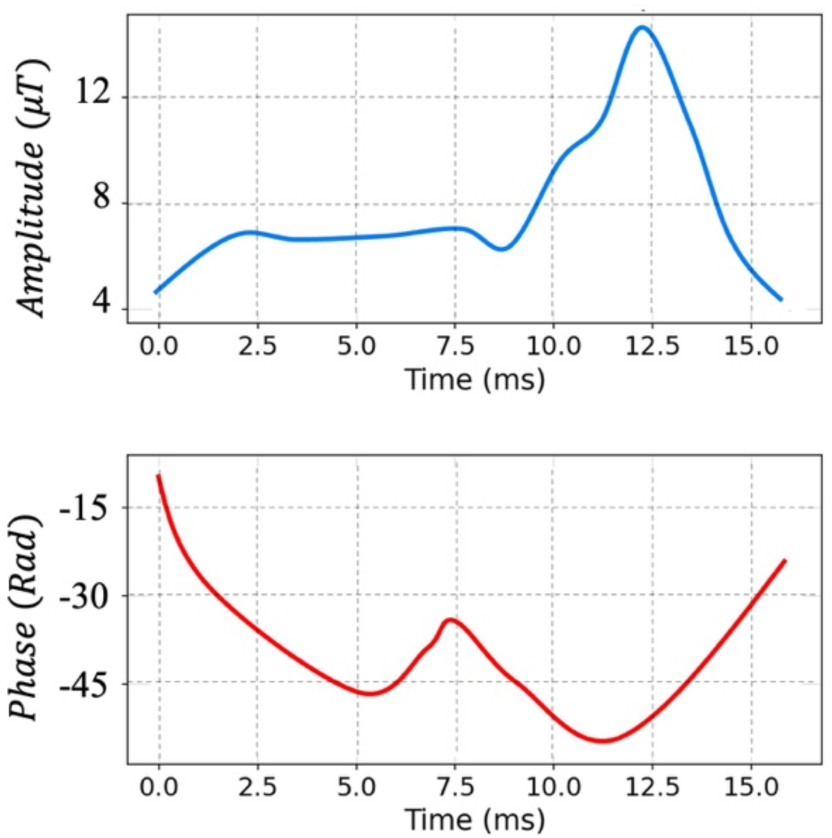
Amplitude and phase waveforms of the self-refocused adiabatic 90°-180° pulse, combining excitation and refocusing into a single pulse.

For timing characterization, (*t_x_*) is defined as the interval between the effective center and the end of the 180° pulse, where the effective center corresponds to the location at which the slope of the applied quadratic phase becomes zero. The resulting self-refocused adiabatic pulse had a total duration of 16 ms. For Δ*T* = 2 *ms* and *t_x_* = 5 *ms*, the effective echo time was TE=2(*t_x_* + Δ*T*)=14 ms. Accordingly, the spin echo was formed 14 ms after the effective center of the 90° excitation component, corresponding to 20 ms from the beginning of the pulse. In comparison, applying the phase-matched 90° and 180° pulses separately, while maintaining a 2-ms interval between the end of the refocusing pulse and signal readout, would result in a minimum TE of 32 ms.

### 2.2. RF Pulse Simulations

Numerical simulations were performed in MATLAB (MathWorks, Natick, MA) to characterize the spectral response and B1 robustness of the FSI self-refocused adiabatic pulse. The simulations were implemented using the discrete-time SLR algorithm, in which the Cayley-Klein parameters, *α* and *β*, were calculated sequentially for the piecewise-constant RF waveform [16]. The resulting parameters were used to determine the transverse magnetization and corresponding magnitude and phase of the pulse spectral response, assuming an initial longitudinal magnetization of unity. This calculation is equivalent to a discrete-time Bloch simulation under the SLR formulation [16]. Relaxation effects were neglected to isolate the intrinsic RF pulse response.

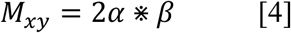

The initial longitudinal magnetization, *M*_0_ is assumed to be 1.

The simulated FSI magnetization profiles demonstrated robust performance over the variation in B1 (Fig. 6). The spectral profiles across B1 scaling factors from 0.8 to 1.2 (Fig. 6a) showed that the passband shape and bandwidth were largely preserved, with B1 variation primarily affecting signal amplitude rather than distorting the spectral response. The corresponding 3D representation (Fig. 6b) further illustrates the stability of the magnetization response as a function of both frequency offset and B_1_ scaling. Peak magnetization retained the nominal response at B_1_ scaling factors of 0.8 and 1.2, respectively, demonstrating the B_1_ robustness of the self-refocused adiabatic FSI pulse. Overall, the simulations demonstrate the trade-off between RF peak amplitude, pulse duration, and spectral-profile robustness when selecting *k* for the self-refocused adiabatic pulse design. An asymmetry was observed between the transition bands, and the passband exhibited a non-flat peak, likely reflecting the nonlinear spectral phase and narrow-band adiabatic design of the self-refocused pulse. The nonuniform passband may introduce frequency-dependent variations in FSI signal intensity, such that spins at different frequency offsets within the passband experience slightly different excitation efficiencies. However, the overall spectral response remained stable across the evaluated B_1_ range.

**Figure 6.**
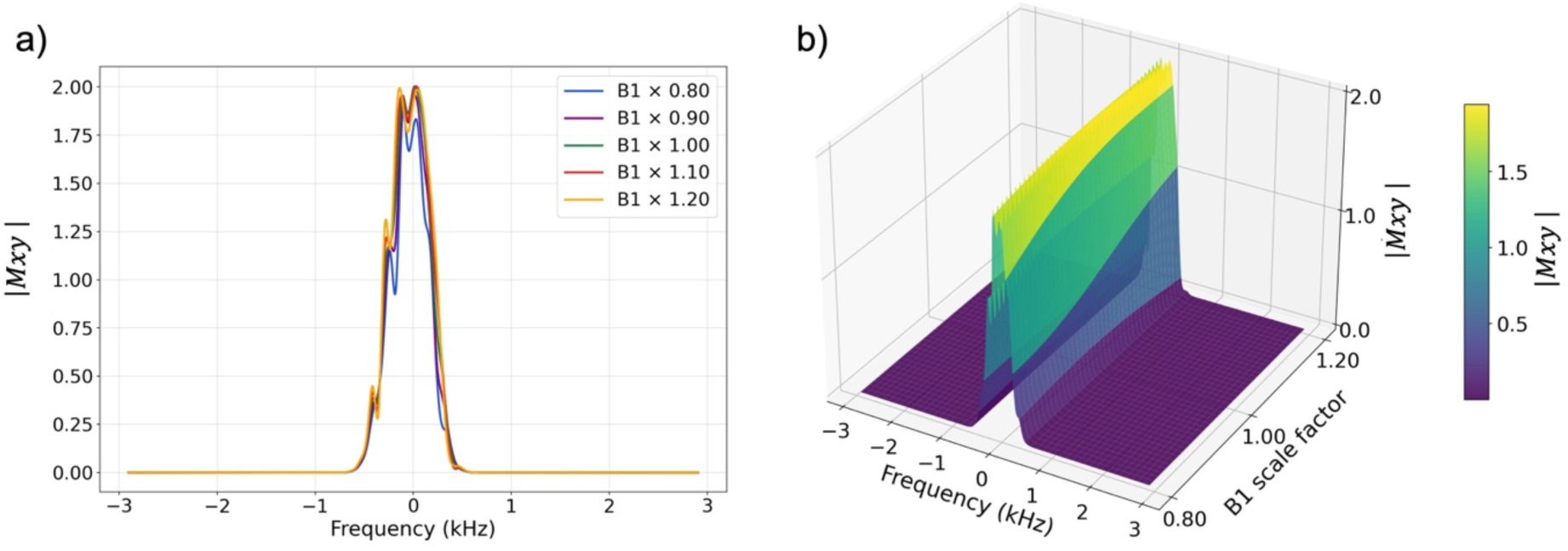
Simulated B1 robustness of the FSI self-refocused adiabatic pulse. a) Magnetization profiles across frequency for varying B1 amplitudes. b) Surface representation of magnetization as a function of frequency offset and B1amplitude, demonstrating the spectral response and B1 insensitivity of the pulse.

### 2.3. Phantom Experiments

Macrophages were cultured in Dulbecco’s Modified Eagle Medium (DMEM) in two T75 flasks. One flask was supplemented with ferumoxytol at a concentration of 30 μg Fe/mL, while the other contained a matched population of unlabeled macrophages serving as the control. Both flasks were incubated for four hours at 37°C to facilitate SPION uptake using a 30 nm average particle size SPION solution (Sigma-Aldrich, St. Louis, MO, USA). Following incubation, all cells were fixed in 4% paraformaldehyde (PFA). Four sample populations were then prepared containing 0%, 25%, 75%, and 100% SPION-labeled cells by volumetric mixing of labeled and unlabeled cell suspensions. Each sample was embedded in 2% agarose gel within individual vials for MR imaging.

All imaging was performed on a 7T whole-body MRI scanner (Magnetom, Siemens Healthineers, Erlangen, Germany) using a single-channel transmit and 32-channel receive head coil (Nova medical). 3D FSI data were acquired at 1.0 mm isotropic resolution, TR/TE= 12/500 ms, with a 140×140 mm^2^ field of view (FOV). FSI images were acquired at multiple frequency offsets by systematically shifting the center frequency of the frequency-selective RF pulse from −800 to +800 Hz in 200-Hz increments.

To establish baseline comparisons, two conventional negative-contrast SPIO imaging techniques were also acquired: (1) 2D SE sequence (TE/TR = 21/2000 ms) which is inherently sensitive to T_2_ effects and susceptibility-induced signal loss; and (2) a 3D gradient echo (GRE) sequence (TE/TR = 5/40ms). The three datasets were compared with respect to SPION localization accuracy, background signal suppression, and positive-contrast performance.

The theoretical basis for quantitative FSI analysis follows the dipole-field model described by Cunningham et al., in which a collection of SPION-labeled cells is approximated as a magnetized sphere (signal intensity) [7, 26]. The local magnetic-field perturbation surrounding the sphere is given by

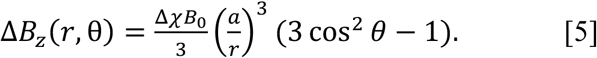

where Δ*χ* is the difference in bulk magnetic susceptibility between the SPION-containing region and its surroundings, *B*_0_ is the main magnetic field, *a* is the radius of the SPION-labeled region, *r* is the distance from its center, and θ is the angle relative to *B*_0_ [26]. This showed that the spatial extent of the frequency-shifted region, and the integrated positive-contrast signal, scales with the volume of labeled cells [26].

To evaluate the relationship between FSI signal intensity and cell concentration, quantitative analysis was performed in MATLAB (MathWorks, Natick, MA, USA). Regions of interest (ROIs) were manually delineated around the hyperintense lobes on nine consecutive image slices for each phantom vial (concentration). ROIs were defined for vials containing 25%, 75%, and 100% labeled cells; the 0% (unlabeled) vial was excluded from quantification as it produced no detectable positive-contrast signal. For each ROI, total signal intensity was computed as the product of the ROI area (in pixels) and the mean signal intensity within the ROI. Measurements were then summed across all nine slices to obtain a representative cumulative signal value for each concentration. Signal intensity ratios between concentrations were subsequently calculated to assess the linearity of the relationship between FSI signal and labeled cell fraction. Imaging was performed with +300 Hz frequency shift.

### 2.4. In Vivo Experiments

In vivo experiments were performed in accordance with the protocol approved by the Icahn School of Medicine at Mount Sinai IACUC. A single adult mouse was used for the in vivo validation study. The mouse was anesthetized via intraperitoneal injection of a ketamine (100 mg/kg) and xylazine (20 mg/kg) solution. Adequate anesthetic depth was confirmed by the absence of the pedal withdrawal reflex prior to proceeding with the injection and imaging. Body temperature was maintained at approximately 37°C using a warm-water circulating pad throughout the imaging session, and respiratory rate was monitored continuously to ensure physiological stability during scanning.

A SPIOs suspension (0.5 ml, 30 nm average particle size; Sigma-Aldrich, St. Louis, MO, USA) was administered via direct intramuscular injection into the hind limb of the mouse. The intramuscular injection route was selected to create a localized depot of SPION at a known anatomical site, enabling direct evaluation of FSI positive-contrast performance against a well-defined region of SPIOs accumulation.

In vivo imaging was performed using the same MRI system and imaging setup as the phantom experiments. Negative-contrast reference images were first acquired using a standard T_2_-weighted SE sequence (TE/TR = 21/2000 ms) to visualize SPION-induced signal voids in the hind limb musculature. Subsequently, 3D FSI data were acquired using the same acquisition protocol with employed in the phantom experiments, +300 Hz frequency shift and 1 mm isotropic resolution, with a total scan time of approximately 11 minutes.

FSI and SE images were compared qualitatively to assess the ability of the FSI technique to convert SPIO-induced signal voids into hyperintense positive-contrast signals at the injection site. The spatial correspondence between regions of signal loss on SE images and regions of signal enhancement on FSI images was evaluated to confirm accurate localization of the SPION depot.

## 3. Results

### 3.1. Phantom experiment

In GRE, SE, and FSI (with zero frequency shifting) images, SPION-containing vials appeared as signal voids, hypointense regions that are difficult to distinguish from other sources of susceptibility-related signal loss (Fig. 7a). In contrast, the FSI with +400 Hz frequency shift composite image depicted the SPION-containing vials as hyperintense positive-contrast signals with improved spatial localization relative to the signal voids observed on conventional sequences. The positive-contrast representation provided by FSI enhanced the conspicuity of the SPION-labeled vials against the suppressed background, facilitating more confident identification of the labeled regions.

**Figure 7.**
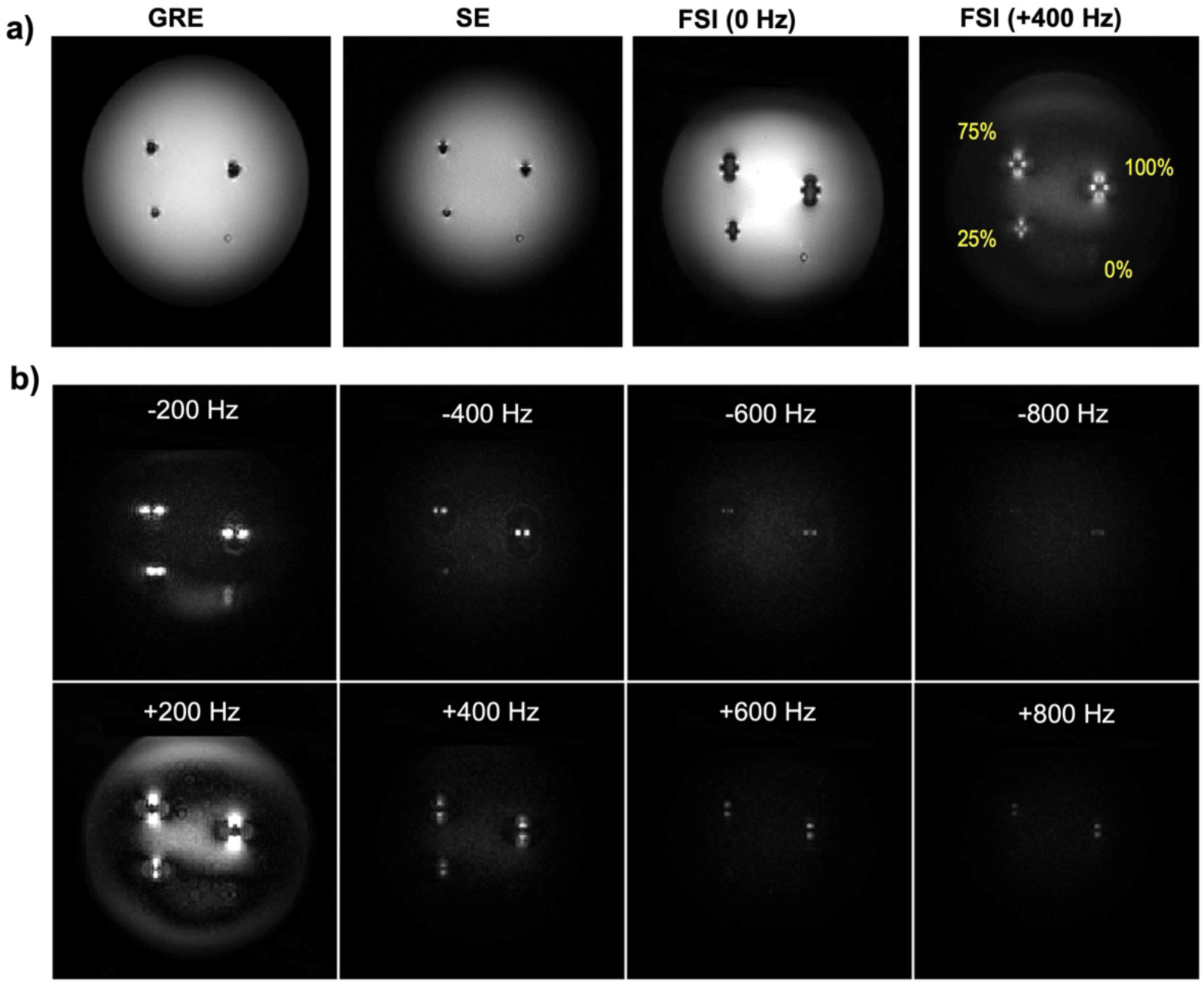
Images obtained from a phantom with SPION labeled cells at 7T MRI. a) Representative images acquired using GRE, SE, FSI without frequency shift (0 Hz), and FSI with a +400 Hz frequency offset, demonstrating differences in SPION-induced contrast among the imaging approaches. b) Images of phantom obtained using the FSI self-refocused pulse. Positive contrast image of phantom using the adiabatic SLR pulse with a spectral offset of -200, -400, -600, and -800 Hz and 200, 400, 600, and 800 Hz.

The magnitude and spatial extent of the excited signal patterns varied systematically with frequency offset. Lower frequency offsets (closer to on-resonance) produced larger excitation patterns with greater signal intensity, reflecting the broader spatial region over which the SPION-induced frequency shift exceeds the excitation threshold (Fig. 7b). Conversely, higher frequency offsets yielded smaller, more spatially confined signal patterns corresponding to regions of stronger local field perturbation in closer proximity to the SPION source. Residual background signal observed at the lowest frequency offsets (±200 Hz) was attributed to incomplete suppression of the on-resonant water peak, a known limitation of narrow-band excitation at offsets [27].

FSI multi-frequency spin echo acquisition revealed characteristic dipole-like signal patterns surrounding the SPION-containing vials. The spatial orientation of the excited regions was frequency-dependent: positive frequency offsets generated hyperintense lobes oriented along the direction of B_0_, while negative frequency offsets produced lobes oriented perpendicular to B_0_. This directional dependence is consistent with the spatial distribution of the dipolar magnetic field perturbation induced by the SPION-containing vials, in which regions of positive frequency shift align parallel to B_0_ and regions of negative frequency shift align in the transverse plane.

Quantitative analysis of the phantom data demonstrated close agreement between the FSI signal intensity and the expected fraction of SPION-labeled cells (Table 1). Signal intensities were normalized to the FSI signal measured in the 100% SPION-labeled cell sample. The normalized signal intensities were 25.02% and 75.16% for samples containing 25% and 75% labeled cells, respectively, demonstrating an approximately proportional relationship between FSI signal intensity and labeled-cell concentration.

**Table 1.** Quantitative FSI signal intensity measurements +300 Hz frequency shift for different SPIO-labeled cell concentrations.

| 100% |  |  |  | 75% |  |  |  | 25% |  |  |  |
| --- | --- | --- | --- | --- | --- | --- | --- | --- | --- | --- | --- |
| Slice | Area | Mean | Signal Intensity | Slice | Area | Mean | Signal Intensity | Slice | Area | Mean | Signal Intensity |
| 1 | 345 | 1385 | 478,050 | 1 | 317 | 1100 | 348,907 | 1 | 118 | 1018 | 120,132 |
| 2 | 323 | 1546 | 499,372 | 2 | 266 | 1227 | 326,458 | 2 | 94 | 996 | 93,666 |
| 3 | 365 | 1503 | 548,665 | 3 | 272 | 1283 | 349,238 | 3 | 92 | 1138 | 104,707 |
| 4 | 333 | 1577 | 525,191 | 4 | 336 | 1196 | 402,125 | 4 | 99 | 1263 | 125,057 |
| 5 | 323 | 1538 | 496,922 | 5 | 275 | 1308 | 359,754 | 5 | 114 | 1218 | 138,913 |
| 6 | 331 | 1442 | 477,612 | 6 | 273 | 1128 | 335,251 | 6 | 90 | 1320 | 118,853 |
| 7 | 258 | 1471 | 379,753 | 7 | 269 | 1191 | 320,441 | 7 | 112 | 1041 | 116,690 |
| 8 | 209 | 1467 | 306,791 | 8 | 237 | 1153 | 273,486 | 8 | 108 | 1029 | 111,182 |
| 9 | 166 | 1153 | 191,461 | 9 | 214 | 1020 | 218,488 | 9 | 57 | 835 | 47,601 |
| Total | n | n | 3,903,821 | Total | n | n | 2,934,152 | Total | n | n | 976,806 |
|  |  | Ratio | 1 |  |  | Ratio | 0.7516 |  |  | Ratio | 0.2502 |

### 3.2 In Vivo imaging

In vivo validation of the FSI technique was performed following intramuscular injection of SPIOs into the hind limb of a mouse model. SE reference images demonstrated characteristic signal voids at the injection site, consistent with the expected T_2_-shortening effects of locally concentrated SPION within the muscle tissue (Fig. 8). These hypointense regions were difficult to delineate precisely against the surrounding musculature due to the inherently low signal contrast between the signal void and adjacent soft tissue structures, a well-recognized limitation of negative-contrast SPION detection. FSI acquisitions of the same anatomical region produced hyperintense positive-contrast signals localized to the SPION injection site, converting the ambiguous signal voids observed on SE imaging into clearly delineated bright regions against a suppressed background. The FSI positive-contrast signal was readily identifiable at the injection site. The enhanced conspicuity of the SPION depot on FSI images relative to SE images demonstrates the practical advantage of positive-contrast detection in living tissue, where multiple endogenous sources of susceptibility variation may otherwise obscure or mimic SPION-induced signal voids. For improved visualization and anatomical orientation, the FSI image was overlaid on a schematic illustration of the mouse to facilitate localization of the SPION-induced positive-contrast signal relative to the injection site.

**Figure 8.**
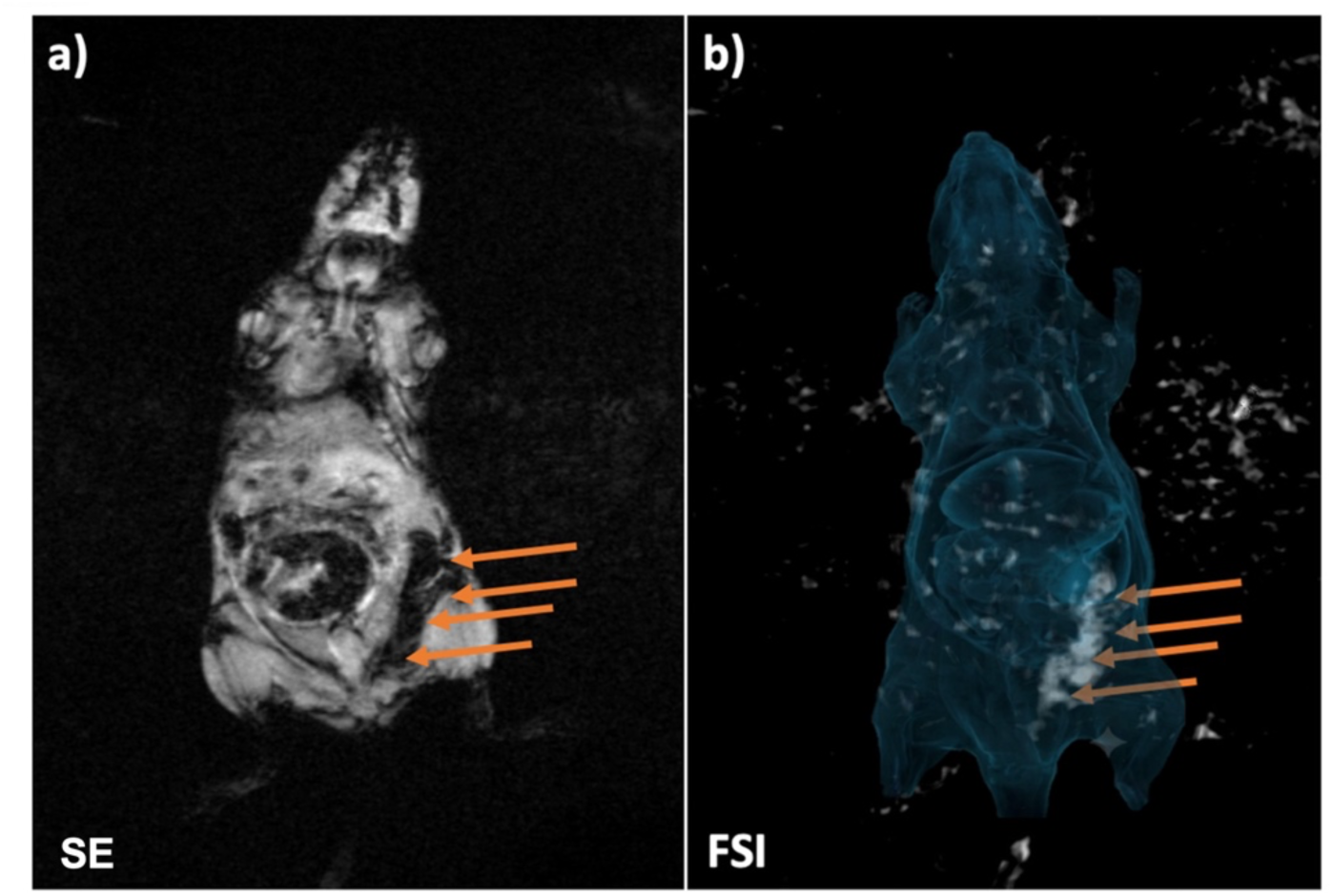
In vivo images of a mouse with SPION injected into the hind legs. Images were obtained at 7T (a) Negative contrast image of mouse acquired using a standard SE sequence. (b) Positive contrast image obtained using FSI with +300 Hz frequency shift. The FSI image was overlaid on a T1-W image to improve visualization of the SPION signal location.

## Discussion and Conclusion

This study introduces FSI, a positive-contrast MRI technique based on a frequency-selective, self-refocused adiabatic SLR pulse for SPION detection at 7T. The pulse-design simulations demonstrated narrow spectral selectivity and robustness to B_1_ variation, phantom experiments demonstrated frequency-dependent positive contrast and an approximately proportional relationship between FSI signal and SPION-labeled cell concentration, and in vivo experiments confirmed the feasibility of detecting SPION in a biological environment. Together, these findings demonstrate the feasibility of FSI as a positive-contrast approach for SPION imaging at ultra-high field.

The FSI pulse was designed to address several challenges associated with SPION imaging at 7T. Although ultra-high-field MRI provides increased SNR and enhanced sensitivity to susceptibility-induced field perturbations [38], pronounced B_0_ and B_1_ inhomogeneities and increased RF power deposition can limit sequence performance [30, 31, 38]. Conventional refocusing pulses are particularly sensitive to B_1_ variation, whereas adiabatic pulses provide greater robustness but generally introduce nonlinear spectral phase and may require additional refocusing pulses, increasing TE and SAR [30–32]. By incorporating adiabatic behavior within a matched-phase, self-refocused SLR framework, FSI combines excitation and refocusing while compensating for nonlinear phase without requiring a second adiabatic 180° pulse. The resulting FSI pulse had a spectral bandwidth of 370 Hz, a peak RF amplitude of 14 μT, and a duration of 16 ms, providing the narrow spectral selectivity required to isolate SPION-induced off-resonant spins. The pulse simulations further demonstrated the effectiveness of the adiabatic design. Across B_1_ scaling factors from 0.8 to 1.2, the spectral response remained largely preserved, with changes in B_1_ primarily affecting signal amplitude rather than substantially shifting or distorting the frequency profile. The simulations also indicated that residual unwanted magnetization was primarily confined to the transition regions, supporting the use of a single self-refocused acquisition in the subsequent phantom and in vivo experiments.

Phantom experiments confirmed that the frequency-selective pulse could translate SPIOs- induced field perturbations into positive contrast. FSI generated characteristic dipole-like hyperintense patterns surrounding SPION-containing vials, with the spatial distribution of the signal changing systematically as the excitation frequency was shifted. In contrast, conventional GRE and SE acquisitions primarily depicted the SPION-containing regions as signal voids. Sampling multiple frequency offsets therefore provided spatial-spectral information about the local susceptibility field while enabling positive visualization of the SPION-containing regions.

An important finding of the phantom study was the relationship between FSI signal intensity and SPION-labeled cell concentration. After normalization to the 100% labeled-cell sample, measured signal intensities were 25.02% and 75.16% for samples containing 25% and 75% labeled cells, respectively. The close agreement between measured signal and expected labeled-cell fractions suggests an approximately proportional relationship under the controlled phantom conditions. Although further validation across a broader range of concentrations is required, these findings indicate that FSI may provide not only positive localization of SPION-labeled cells but also a potential basis for quantitative estimation of labeled-cell burden.

The in vivo experiments further demonstrated that FSI can generate positive contrast in a heterogeneous biological environment. At the intramuscular SPION injection site, conventional imaging produced a hypointense signal void, whereas frequency-shifted FSI acquisitions generated localized hyperintense signal with the characteristic dipolar spatial pattern observed in the phantom experiments. The preservation of this frequency-dependent pattern in vivo supports the interpretation that the detected signal originates from the local SPION-induced field perturbation and demonstrates translation of the technique from controlled phantom conditions to biological tissue.

Two aspects warrant further investigation. First, the in vivo experiments were designed as a proof-of-concept using localized intramuscular SPION injection in a single animal. Future studies should extend FSI to longitudinal cell-tracking models with more heterogeneous distributions of SPION-labeled cells and evaluate its detection sensitivity at lower concentrations. Second, although the phantom experiments demonstrated an approximately proportional relationship between FSI signal and labeled-cell concentration, quantitative performance remains to be validated in vivo, where tissue heterogeneity, B_1_ variations, physiological motion, and endogenous susceptibility effects may influence the measured signal. Future studies incorporating histological correlation will help establish the relationship between FSI signal intensity and local SPION-labeled cell concentration. Future work will include a systematic comparison of the same FSI positive-contrast sequence at 1.5T, 3T, and 7T to characterize the effects of field strength on SPIO detection sensitivity, frequency-shift behavior, image contrast, and overall sequence performance.

## Notes

### Competing Interest Statement

The authors have declared no competing interest.

## References

1. Li, H. and T.J. Meade, Molecular Magnetic Resonance Imaging with Gd(III)-Based Contrast Agents: Challenges and Key Advances. J Am Chem Soc, 2019. 141(43): p. 17025–17041.

2. Wahsner, J., et al., Chemistry of MRI Contrast Agents: Current Challenges and New Frontiers. Chem Rev, 2019. 119(2): p. 957–1057.

3. Lu, Z.R., V. Laney, and Y. Li, Targeted Contrast Agents for Magnetic Resonance Molecular Imaging of Cancer. Acc Chem Res, 2022. 55(19): p. 2833–2847.

4. Bulte, J.W. and D.L. Kraitchman, Iron oxide MR contrast agents for molecular and cellular imaging. NMR Biomed, 2004. 17(7): p. 484–99.

5. Wang, Y.X., et al., Recent advances in superparamagnetic iron oxide nanoparticles for cellular imaging and targeted therapy research. Curr Pharm Des, 2013. 19(37): p. 6575–93.

6. Rosen, J.E., et al., Iron oxide nanoparticles for targeted cancer imaging and diagnostics. Nanomedicine, 2012. 8(3): p. 275–90.

7. Bakhtiary, Z., et al., Targeted superparamagnetic iron oxide nanoparticles for early detection of cancer: Possibilities and challenges. Nanomedicine, 2016. 12(2): p. 287–307.

8. Bulte, J.W.M., C. Wang, and A. Shakeri-Zadeh, In Vivo Cellular Magnetic Imaging: Labeled vs. Unlabeled Cells. Adv Funct Mater, 2022. 32(50).

9. Neuwelt, A., et al., Iron-based superparamagnetic nanoparticle contrast agents for MRI of infection and inflammation. AJR Am J Roentgenol, 2015. 204(3): p. W302–13.

10. Jeon, M., et al., Iron Oxide Nanoparticles as T(1) Contrast Agents for Magnetic Resonance Imaging: Fundamentals, Challenges, Applications, and Prospectives. Adv Mater, 2021. 33(23): p. e1906539.

11. Wei, H., et al., Exceedingly small iron oxide nanoparticles as positive MRI contrast agents. Proc Natl Acad Sci U S A, 2017. 114(9): p. 2325–2330.

12. Corot, C., et al., Recent advances in iron oxide nanocrystal technology for medical imaging. Adv Drug Deliv Rev, 2006. 58(14): p. 1471–504.

13. Rümenapp, C., B. Gleich, and A. Haase, Magnetic nanoparticles in magnetic resonance imaging and diagnostics. Pharm Res, 2012. 29(5): p. 1165–79.

14. Wang, X., et al., Application of nanotechnology in cancer therapy and imaging. CA Cancer J Clin, 2008. 58(2): p. 97–110.

15. Liu, T., et al., Unambiguous identification of superparamagnetic iron oxide particles through quantitative susceptibility mapping of the nonlinear response to magnetic fields. Magn Reson Imaging, 2010. 28(9): p. 1383–9.

16. Liu, W., et al., In vivo MRI using positive-contrast techniques in detection of cells labeled with superparamagnetic iron oxide nanoparticles. NMR Biomed, 2008. 21(3): p. 242–50.

17. Jung, H., et al., Dual MRI T1 and T2(*) contrast with size-controlled iron oxide nanoparticles. Nanomedicine, 2014. 10(8): p. 1679–89.

18. Reimer, P. and B. Tombach, Hepatic MRI with SPIO: detection and characterization of focal liver lesions. Eur Radiol, 1998. 8(7): p. 1198–204.

19. Deng, L.H., et al., Size and PEG Length-Controlled PEGylated Monocrystalline Superparamagnetic Iron Oxide Nanocomposite for MRI Contrast Agent. Int J Nanomedicine, 2021. 16: p. 201–211.

20. Alipour, A., et al., A new class of cubic SPIONs as a dual-mode T1 and T2 contrast agent for MRI. Magn Reson Imaging, 2018. 49: p. 16–24.

21. Sharma, V.K., et al., Highly monodisperse low-magnetization magnetite nanocubes as simultaneous T(1)-T(2) MRI contrast agents. Nanoscale, 2015. 7(23): p. 10519–26.

22. Mani, V., et al., Gradient echo acquisition for superparamagnetic particles with positive contrast (GRASP): sequence characterization in membrane and glass superparamagnetic iron oxide phantoms at 1.5T and 3T. Magn Reson Med, 2006. 55(1): p. 126–35.

23. Stuber, M., et al., Positive contrast visualization of iron oxide-labeled stem cells using inversion-recovery with ON-resonant water suppression (IRON). Magn Reson Med, 2007. 58(5): p. 1072–7.

24. Zhao, Q., et al., Positive contrast technique for the detection and quantification of superparamagnetic iron oxide nanoparticles in MRI. NMR Biomed, 2011. 24(5): p. 464–72.

25. Diwoky, C., et al., Positive contrast of SPIO-labeled cells by off-resonant reconstruction of 3D radial half-echo bSSFP. NMR Biomed, 2015. 28(1): p. 79–88.

26. Cunningham, C.H., et al., Positive contrast magnetic resonance imaging of cells labeled with magnetic nanoparticles. Magn Reson Med, 2005. 53(5): p. 999–1005.

27. Balchandani, P., et al., Self-refocused spatial-spectral pulse for positive contrast imaging of cells labeled with SPIO nanoparticles. Magn Reson Med, 2009. 62(1): p. 183–92.

28. Zarghami, N., et al., Optimization of molecularly targeted MRI in the brain: empirical comparison of sequences and particles. Int J Nanomedicine, 2018. 13: p. 4345–4359.

29. Shen, Y., et al., Detecting sub-voxel microvasculature with USPIO-enhanced susceptibility-weighted MRI at 7 T. Magn Reson Imaging, 2020. 67: p. 90–100.

30. Balchandani, P., et al., Self-refocused adiabatic pulse for spin echo imaging at 7 T. Magn Reson Med, 2012. 67(4): p. 1077–85.

31. van Kalleveen, I.M., et al., Adiabatic turbo spin echo in human applications at 7 T. Magn Reson Med, 2012. 68(2): p. 580–7.

32. Pauly, J., et al., Parameter relations for the Shinnar-Le Roux selective excitation pulse design algorithm [NMR imaging]. IEEE Trans Med Imaging, 1991. 10(1): p. 53–65.

33. Balchandani, P., J. Pauly, and D. Spielman, Designing adiabatic radio frequency pulses using the Shinnar-Le Roux algorithm. Magn Reson Med, 2010. 64(3): p. 843–51.

34. Feldman, R.E. and P. Balchandani, A semiadiabatic spectral-spatial spectroscopic imaging (SASSI) sequence for improved high-field MR spectroscopic imaging. Magn Reson Med, 2016. 76(4): p. 1071–82.

35. Ordidge, R., et al., Ultrahigh field brain magnetic resonance imaging using semiadiabatic radiofrequency pulses. NMR Biomed, 2022. 35(6): p. e4672.

36. Feldman, R.E., et al., A SEmi-Adiabatic matched-phase spin echo (SEAMS) PINS pulse-pair for B1 -insensitive simultaneous multislice imaging. Magn Reson Med, 2016. 75(2): p. 709–17.

37. Balchandani, P. and D. Qiu, Semi-adiabatic Shinnar-Le Roux pulses and their application to diffusion tensor imaging of humans at 7T. Magn Reson Imaging, 2014. 32(7): p. 804–12.

38. Opheim, G., et al., 7T Epilepsy Task Force Consensus Recommendations on the Use of 7T MRI in Clinical Practice. Neurology, 2021. 96(7): p. 327–341.

